# pDeep3: Towards More Accurate Spectrum Prediction with Fast Few-Shot Learning

**DOI:** 10.1101/2020.09.13.295105

**Authors:** Ching Tarn, Wen-Feng Zeng

## Abstract

Spectrum prediction using deep learning has attracted a lot of attention in recent years. Although existing deep learning methods have dramatically increased the prediction accuracy, there is still considerable space for improvement, which is presently limited by the difference of fragmentation types or instrument settings. In this work, we use the few-shot learning method to fit the data online to make up for the shortcoming. The method is evaluated using ten datasets, where the instruments includes Velos, QE, Lumos, and Sciex, with collision energies being differently set. Experimental results show that few-shot learning can achieve higher prediction accuracy with almost negligible computing resources. For example, on the dataset from a untrained instrument Sciex-6600, within about 10 seconds, the prediction accuracy is increased from 69.7% to 86.4%; on the CID (collision-induced dissociation) dataset, the prediction accuracy of the model trained by HCD (higher energy collision dissociation) spectra is increased from 48.0% to 83.9%. It is also shown that, the method is not critical to data quality and is sufficiently efficient to fill the accuracy gap. The source code of pDeep3 is available at http://pfind.ict.ac.cn/software/pdeep3.

---

Machine learning algorithms currently play important roles in mass spectrometry-based proteomics.^1–4^ In recent years, deep learning-based methods have attracted more and more attention, for both data-dependent acquisition (DDA) and data-independent acquisition (DIA) data analysis.^5– 16^

For DDA data analysis, the recently published pDeep,^5^ Prosit,^6^ and DeepMass:Prism^7^ showed that integrating spectrum prediction into DDA search engines would dramatically increase the identification rates. Our recent work, pValid,^17^ further demonstrated that the spectra predicted by pDeep could be applied to filter out unreliable peptide spectrum matches (PSMs), resulting in more accurate peptide identifications. We also used the predicted spectra to re-rank the *de novo* sequenced peptides to achieve higher recall and precision.^18^

For DIA data analysis, Prosit and DeepMass:Prism showed that the performance of predicted spectral libraries was close to that of experimental ones in spectral library search. DeepDIA built predicted spectral libraries of peptides from nearly the full human proteome for DIA analysis and detected more proteins than those using the pre-built DDA spectral libraries.^11^ Searle *et al*. also used a full proteome library predicted by Prosit and corrected it using empirical spectra.^19^ A recent work combined the experimental spectral library with the predicted library for proteins of interest to deepen the coverage of targeted protein families.^20^

It turns out that more accurate predictions would result in better identification performance for both DDA and DIA analysis.^6,7^ However, in practical uses, not all MS/MS predictions are sufficiently accurate.^9,11,21^ Prosit tried to correct the predicted spectra by enumerating all predicting collision energies; DeepDIA aimed at training instrument-specific models for more accurate predictions. Unfortunately, either enumerating all energies or training models for specific instruments is time-consuming, and hence, accelerating devices such as GPUs are indispensable. Moreover, our previous work, pDeep2, showed that transfer learning is an effective way to improve the prediction of modified peptides from a pre-trained unmodified model requiring only tens of thousands of spectra with modifications for training. Due to the computational costs, the training step could be done within minutes using a GPU, but it will take tens of minutes or even hours when only CPUs are available.

Pre-trained offline models can hardly achieve the same accuracy as online-trained models since the latter can automatically learn the characteristics of different MS fragmentation and instrument settings. However, online learning from scratch requires many computational resources to process spectra in real-time and needs a lot of data from the specific instrument. Inspired by the few-shot learning technique,^22^ we use a few-shot fine-tuning method to train models for specific instruments based on a pre-trained offline model using only hundreds of training spectra, achieving higher accuracy. As the name “few-shot” implies, few-shot learning refers to the practice of learning a model with a small amount of training data and has been widely used in computer vision and other machine learning tasks. There are two advantages of few-shot learning (also called fine-tuning in this work): 1) fine-tuning a low accuracy pre-trained model would increase the accuracy of predictions; 2) fine-tuning could be done within only about 10 seconds on CPU, making training lab-specific or even sample-specific models applicable on a laptop or a personal computer. The source code of pDeep3 is available at http://pfind.ict.ac.cn/software/pdeep3.

## Materials and Methods

### Data preparation

Diverse datasets are collected from the test datasets of our previous work,^9^ which are all publicly available.^23–27^ For each dataset, we only consider spectra from a subset of RAW files, which are sufficient for fine-tuning and testing. Dataset information is listed in Table Sciex-6600 TripleTOF-based CID (Collision-Induced Dissociation) data and LIT (Linear Ion Trap)-based CID data are also considered. For LIT-based CID data in ProteomeTools datasets, MaxQuant search results at 0.1% FDR are used, and b/y ions are extracted within ± 0.5 Da matching mass tolerance.

### Spectrum similarity measurement

We use Pearson correlation coefficient (PCC) to measure the similarity between the predicted and experimental spectra. For a given spectrum and its identified peptide, the intensities of all b_i_ and y_i_ ions with 1+ and 2+ charge states are extracted from the spectrum, where i = 1, 2, …, *l* − 1, and *l* is the peptide length. This 4 × (*l* − 1) intensity matrix is then reshaped into a 1-dim vector *x*. The predicted 4 × (*l* − 1) b/y intensity matrix is reshaped into a 1-dim vector *y*, then PCC can be calculated as

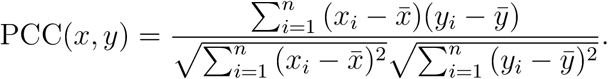

We used the median PCC, PCC_>0.75_ and PCC_>0.90_ to evaluate the performance of different methods, where PCC_>*t*_ is defined as the proportion of PCCs greater than *t* in the corresponding dataset.

### The few-shot learning procedure

The few-shot learning procedure of pDeep3 is illustrated in Figure 1. The pre-trained model was trained using millions of spectra from different instruments and collision energies, as described in our previous work.^9^ For a specific dataset, the network parameters could be then fine-tuned with only a few hundred spectra to achieve better predictions. The main hyper-parameters of fine-tuning are: epoch number = 2, learning rate = 0.001, mini-batch size = 1024. Since the peptides with different lengths are assigned into different mini-batch, the actual size of each mini-batch depends on peptides’ length distribution.

**Figure 1:**
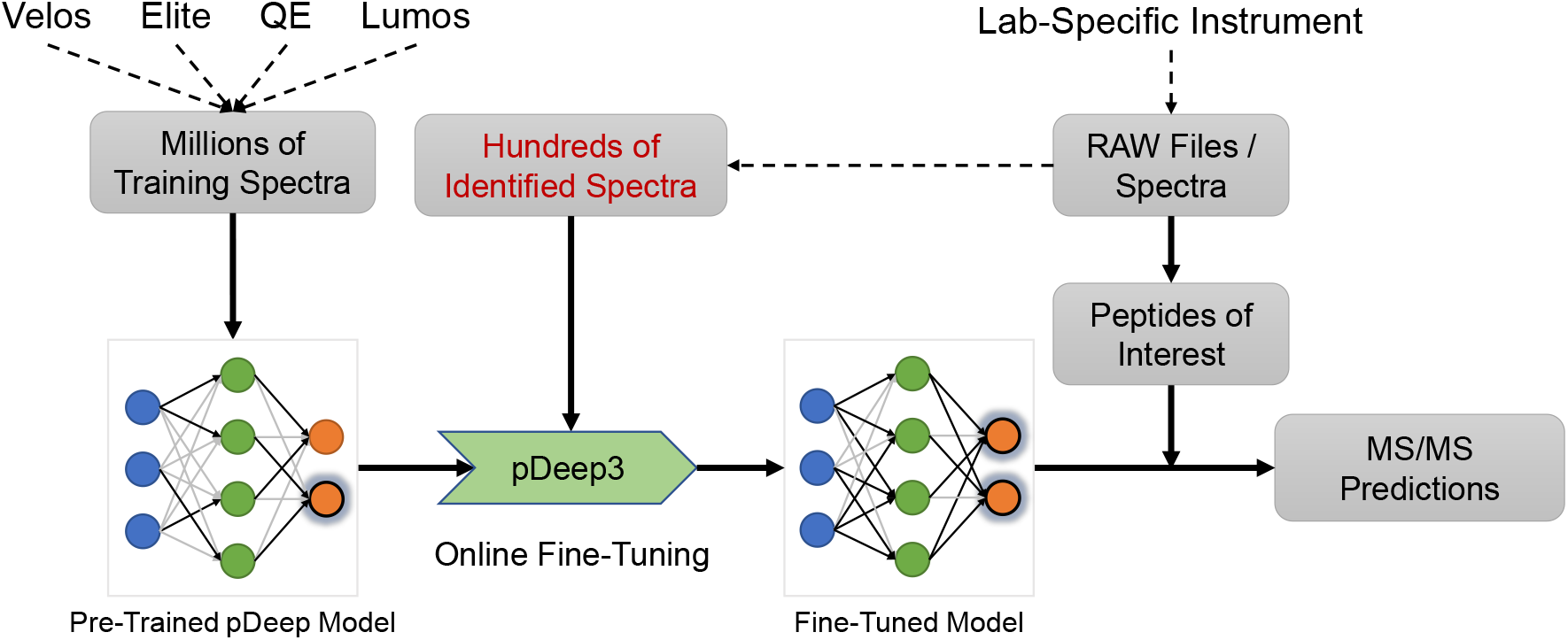
Workflow of the fine-tuning procedure of pDeep3.

The quality of the pre-trained model is the key to fast fine-tuning. It ensures that the network parameters are already close to the optimal ones for a new dataset, and hence we can tune the model to fit the new spectra with only a few back-propagation updates. As reported in our previous work,^9^ the pre-trained pDeep model was trained and tested on Velos, Elite, QE, and Lumos datasets. It achieved about 0.98 median PCCs in most test datasets, indicating that the pre-trained model is suitable for fine-tuning. ^*^

### Grid search of parameters for unseen instruments

In the current version of pDeep3, the instrument type is encoded as a 5-dimensional (5-dim) one-hot vector, corresponding to QE (including QE, QE+, QE-HF, and QE-HFX), Velos, Elite, Fusion, and Lumos, respectively. When the instrument type is not in the above instrument list, pDeep3 could use the grid search strategy to find a suitable instrument and NCE parameter combination for a given training dataset of this unseen instrument. For the grid search, pDeep3 enumerates all combinations of the above five instrument types and the NCE values ranging from 10 to 45, and then determines the instrument and NCE by the best PCC_>0.90_. Fine-tuning is further applied based on the determined instrument and NCE to obtain a better model.

## Results and Discussions

### Hyper-parameter selection

In the fine-tuning step, the number of epochs *e* and the size of the training set *n* are two key hyper-parameters for the trade-off of the training cost and prediction accuracy. Here we use three sets of experiments to select these hyper-parameters. It is worth mentioning that, for each dataset, we only partially select peptide-spectrum matches (PSMs) from a RAW file when fine-tuning, and test the fine-tuned model using all PSMs from another RAW file.

To select an appropriate *e* in the fine-tuning step, we try *e* in {0, 1, …, 8}, and set *n* as 100. The test results are shown in Figure 2a. It turns out that, as *e* increases, the performance on most of the datasets improves as well, and the test PCC_>0.75_ and PCC_>0.90_ become stable after about two epochs. In particular, as the pre-trained model has been trained by other peptides of ProteomeTools,^9^ the test results on PL are already close to the best accuracy, and hence there is no further improvement in the PL dataset as *e* increases (Figure 2a). However, the performance is not decreasing, indicating that fine-tuning may not suffer from the potential risk of over-fitting.

**Figure 2:**
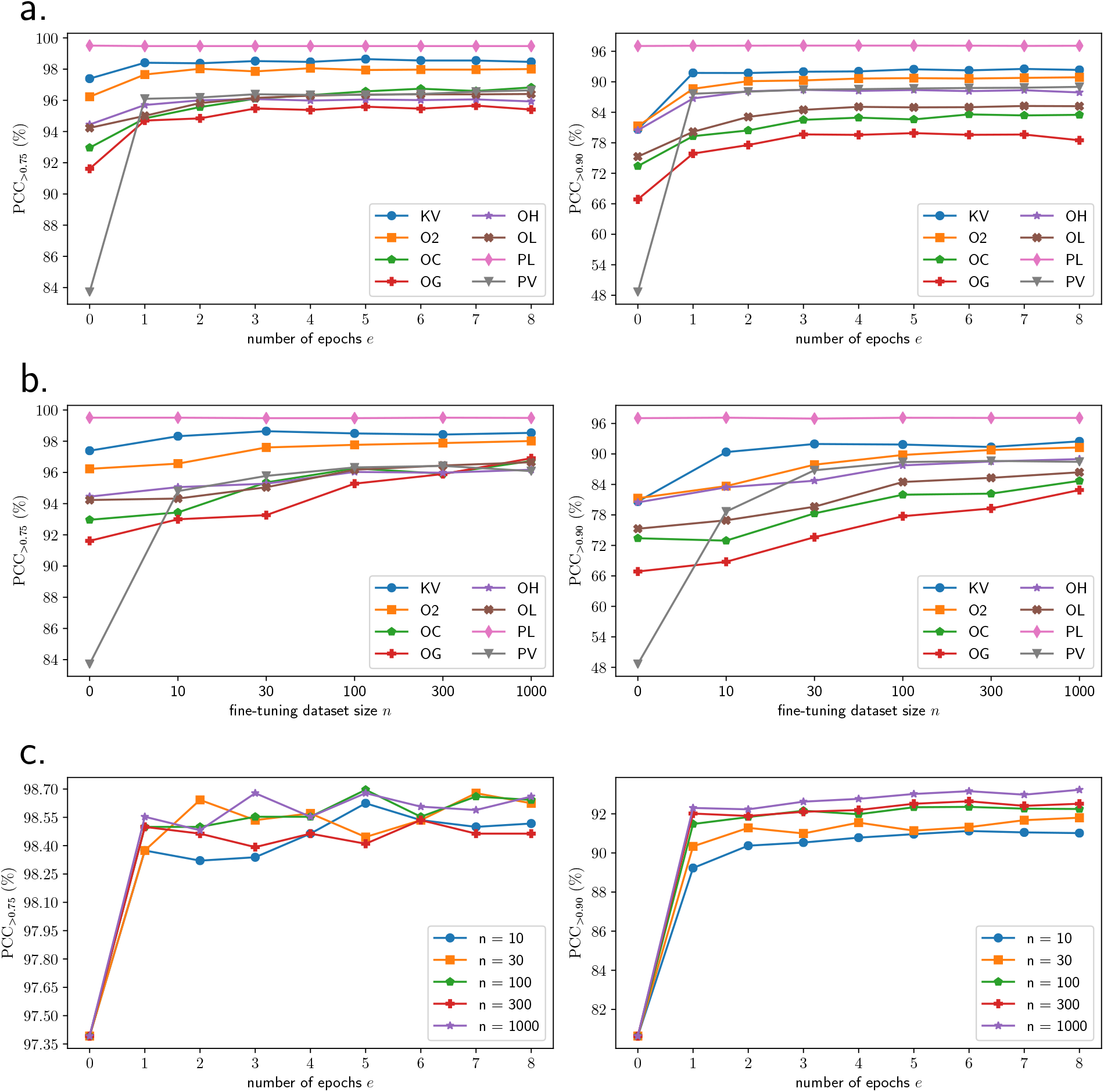
Tests of different hyper-parameters, training epochs *e* and training set size *n*, on different datasets. (a) Tests of *e* with *n* = 100. (b) Tests of *n* with *e* = 2. (c) Tests of different combinations of *e* and *n* on the KV dataset.

We also try *n* in {0, 10, 30, 100, 300, 1000}, and set *e* as 2 for fine-tuning. The results are demonstrated in Figure 2b. The performance steadily gets better as *n* increases. And for PL, the results are not much changed as well. The improvements on most datasets gradually slow down as *n* increases after *n* > 100. Specifically, the improvements on KV have achieved the optimal when only about 10 or 30 samples are used; for Olsen’s datasets (O2, OC, OG, OH, OL), the increases are roughly linearly related to log *n*.

We further try different combinations of *e* and *n* for *e* ∈ {0, 1, …, 8} and *n* ∈ {10, 30, 100, 300, 1000}. The test results on the KV dataset are illustrated in Figure 2c. As shown in Figure 2c, the effects of *e* and *n* are quite independent. That means even if *n* is different, the method achieves a large increase after about two epochs, and the improvement slows down thereafter. With different numbers of epochs *e*, the larger the size of the training set, the better the results could be, and the growth rate gradually slows down as *n* increases. Results on other datasets can be found in Data S1.

As the time required for fine-tuning is relative to the number of epochs *e* and the size of the training set *n*, to ensure that fine-tuning can be done using CPU within the acceptable time, it is necessary to limit the scale without significantly sacrificing the performance. The test results in Figure 2 demonstrate that the fine-tuned model can reach good performance in both fine-tuning speed and predicting accuracy using 100 training samples and two epochs. In these tests, the fine-tuning can be finished within only about 10 seconds using an Intel CPU (Xeon CPU E5-2680 v4 @ 2.40GHz, without GPU devices), showing that fine-tuning is always recommended before predicting for any datasets.

### Intra-RAW and inter-RAW tests

For a dataset, we question whether we need to fine-tune different models for different RAW files of the dataset (run-specific fine-tuning), or we only need to fine-tune a model for all RAW files (sample-/instrument-specific fine-tuning). To answer it, we apply intra-RAW and inter-RAW test experiments.

For each dataset, we split PSMs in each RAW file into a training set (100 PSMs) and a testing set (the rest of the RAW). The pre-trained model is fine-tuned by 5 training sets, respectively (*e* = 2, *n* = 100), resulting in 5 fine-tuned models. Each model is then tested separately on all the five testing sets, and we get a 5 × 5 PCC_>*x*_ matrix S. Each set of results of a model corresponds to a row of the matrix S, where S_*ij*_ is the result of fine-tuning with RAW file *i* and testing with RAW file *j*. Test results on the five RAW files of KV and O2 datasets are listed in Table 2. We use the standard deviation (σ) to measure the similarity between the results. On the same testing RAW file, fine-tuning by the training set of any RAW file achieves the similar PCC_>0.75_ (σ = 0.56% for KV and σ = 0.17% for O2) and PCC_>0.90_ (σ = 1.51% for KV and σ = 0.38% for O2), indicating that the model can be fine-tuned only once on a RAW file and applies to others in the same dataset if the instrument settings are not changed. Results of other datasets can be found at Data S2.

**Table 1:** Dataset information

| Dataset | Description | Abbr. | Instrument | NCE | #RAWs | #PSMs |
| --- | --- | --- | --- | --- | --- | --- |
| Kuster-Velos | Human, Milk | KV | Velos | 40 | 5 | 35740 |
| Olsen-293-QE | HEK293T, Trypsin | O2 | QE | 28 | 5 | 71788 |
| Olsen-HeLa-QE | HeLa, Trypsin | OH | QE | 28 | 5 | 106006 |
| Olsen-GC-QE | HeLa, GluC | OG | QE | 28 | 5 | 74826 |
| Olsen-CT-QE | HeLa, Chymotrypsin | OC | QE | 28 | 5 | 60191 |
| Olsen-LC-QE | HeLa, LysC | OL | QE | 28 | 5 | 98985 |
| PT-Lumos | ProteomeTools | PL | Lumos | 30 | 5 | 41772 |
| Pandey-Velos | Human, Fetal-Gut | PV | Velos | 35 | 5 | 46114 |
| LFQBench-HE | HeLa and E.coli | - | Sciex-6600 | - | 2 | 28333 |
| PT-CID | ProteomeTools | - | Lumos | 35 | 2 | 12345 |

**Table 2:** Intra-RAW and inter-RAW tests. *r*_*i*_ means the model is trained on the training set from RAW *i*, and *s*_*j*_ means it is tested on the testing set from RAW file *j*.

| (a) KV |  |  |  |  |  |  |  |  |  |  |
| --- | --- | --- | --- | --- | --- | --- | --- | --- | --- | --- |
| | $\text{PCC}_{>0.75}$ ($\%$, $\sigma = 0.56\%$) | | | | | $\text{PCC}_{>0.90}$ ($\%$, $\sigma = 1.51\%$) | | | | |
| | $s_1$ | $s_2$ | $s_3$ | $s_4$ | $s_5$ | $s_1$ | $s_2$ | $s_3$ | $s_4$ | $s_5$ |
| $r_1$ | 98.65 | 98.34 | 98.66 | 97.40 | 99.21 | 93.21 | 91.84 | 94.26 | 92.69 | 95.84 |
| $r_2$ | 98.76 | 98.69 | 98.75 | 97.74 | 99.27 | 93.10 | 91.65 | 93.32 | 92.46 | 96.10 |
| $r_3$ | 98.65 | 98.42 | 98.66 | 97.57 | 99.23 | 93.25 | 91.74 | 93.75 | 92.57 | 95.98 |
| $r_4$ | 98.78 | 98.53 | 98.73 | 97.62 | 99.21 | 94.25 | 92.17 | 93.89 | 93.06 | 96.22 |
| $r_5$ | 98.57 | 98.48 | 98.66 | 97.50 | 99.21 | 92.88 | 91.11 | 93.05 | 92.19 | 95.86 |

| (b) O2 |  |  |  |  |  |  |  |  |  |  |
| --- | --- | --- | --- | --- | --- | --- | --- | --- | --- | --- |
| | $\text{PCC}_{>0.75}$ ($\%$, $\sigma = 0.17\%$) | | | | | $\text{PCC}_{>0.90}$ ($\%$, $\sigma = 0.38\%$) | | | | |
| | $s_1$ | $s_2$ | $s_3$ | $s_4$ | $s_5$ | $s_1$ | $s_2$ | $s_3$ | $s_4$ | $s_5$ |
| $r_1$ | 97.64 | 97.89 | 97.54 | 97.92 | 97.89 | 89.70 | 90.12 | 89.40 | 89.34 | 89.24 |
| $r_2$ | 97.74 | 98.00 | 97.61 | 97.96 | 98.01 | 90.04 | 90.44 | 89.52 | 89.73 | 89.91 |
| $r_3$ | 97.74 | 97.98 | 97.70 | 98.01 | 97.99 | 89.92 | 90.14 | 89.97 | 89.57 | 89.78 |
| $r_4$ | 97.50 | 97.85 | 97.55 | 97.90 | 97.89 | 90.00 | 90.43 | 89.69 | 90.27 | 90.13 |
| $r_5$ | 97.62 | 97.88 | 97.56 | 97.95 | 97.91 | 89.27 | 89.50 | 89.07 | 89.40 | 89.66 |

### Tests of training set selection bias

One concern of the fine-tuning step is that the 100 randomly selected PSMs hardly cover all peptide characteristics (different lengths, different charge states, etc.). Once some properties of peptides are not selected for fine-tuning, the fine-tuned model may prefer the selected properties, resulting in poor predictions on peptides with non-selected properties. We call it “training set selection bias” of the fine-tuning model. To test whether the bias exists, we evaluate six training set selection methods, as listed in follows:

1. *low*. The training set only contains peptides with precursor charge ≤ 3;
2. *high*. The training set only contains peptides with precursor charge > 3;
3. *short*. The training set only contains peptides with length ≤ 12;
4. *long*. The training set only contains peptides with length > 12;
5. *mod*. The training set only contains peptides with Oxidation[M];
6. *unmod*. The training set only contains peptides without any variable modification.

We also consider “random selection” (denoted as *random*) and “no fine-tuning” (denoted as *none*) for comparisons. All the models are fine-tuned by selected training sets (*e* = 2, *n* = 100) and tested by another RAW file. The results is illustrated in Figure 3, and show that, in most cases, a model with fine-tuning is always better than that without fine-tuning. Random selection usually achieves the best performance.

**Figure 3:**
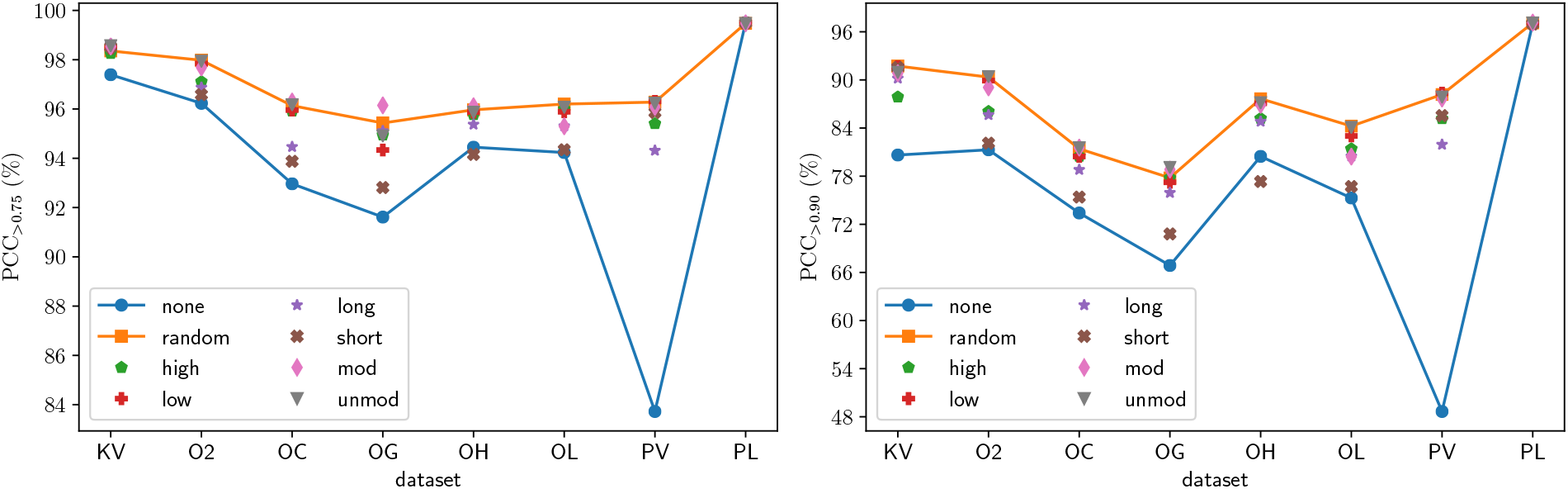
Test of different training set selection methods for fine-tuning.

### Grid search and fine-tuning on Sciex-6600 and LIT-based CID datasets

Although the TripleTOF-based CID of Sciex-6600 is also the beam-type CID which is the same as Orbitrap-based HCD, there are still many differences between Sciex-6600 and Orbitrap-based instruments, including the collision energy settings, the mass analyzer, ion detector, signal processing, etc. As the pDeep model is only trained on HCD data, we use grid search to find the best instrument and NCE for the Sciex-6600 dataset before fine-tuning (Elite@NCE=31, as shown in Figure 4a). Even after the best instrument and NCE are determined, fine-tuning is still very useful to obtain a better model. As shown in Figure 4c, PCC_>0.90_ increases from 69.7% to 86.4% after fine-tuning with *n* = 100 and *e* = 8.

**Figure 4:**
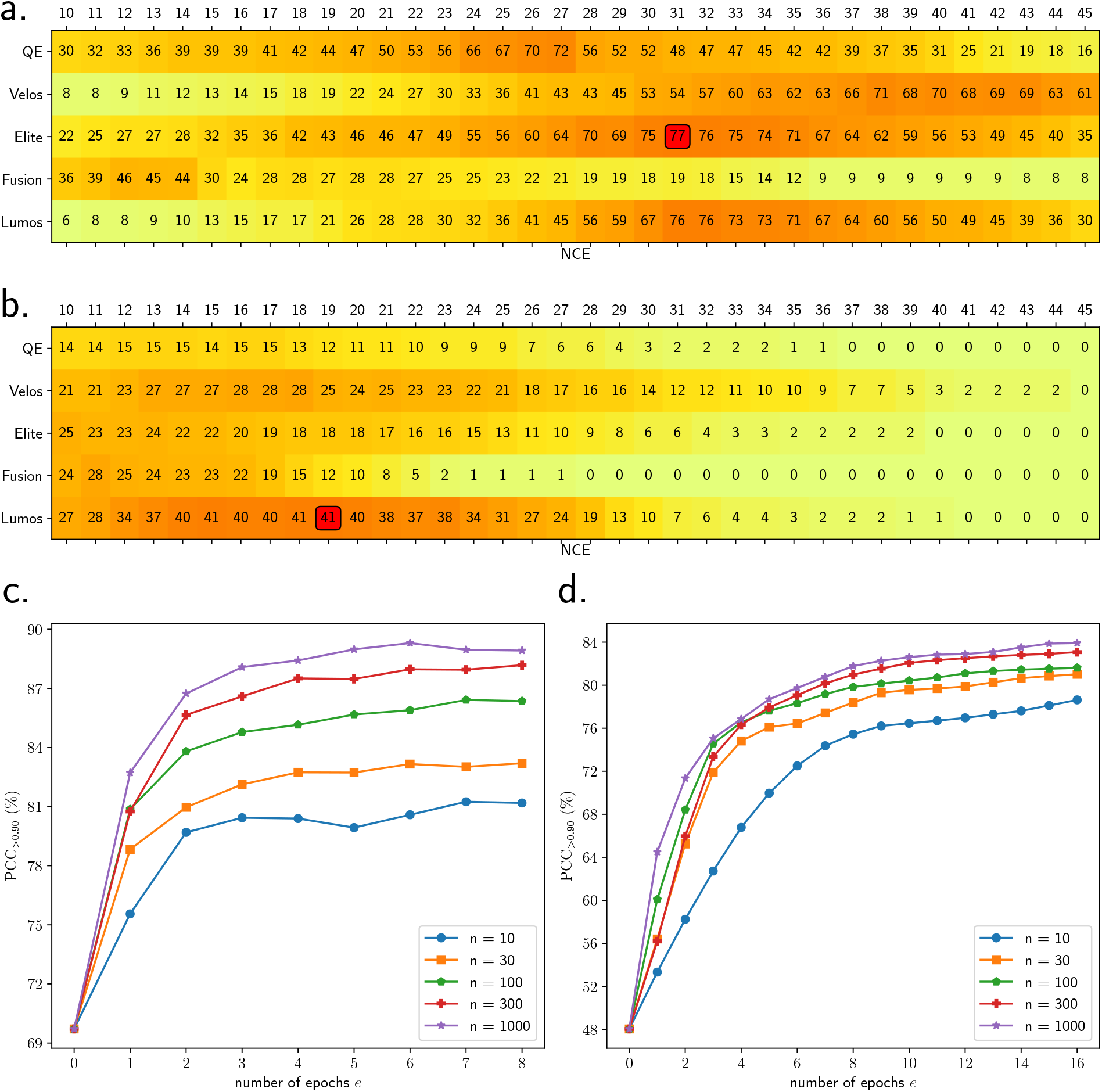
Grid search and few-shot learning for Sciex-6600 instrument and LIT-based CID spectra. (a) PCC_>0.90_ of each setting of grid search for best instrument and NCE for Sciex-6600 data. The best setting is Elite@NCE=31. The grid search takes 59.4 seconds. (b) PCC_>0.90_ of each setting of grid search for best instrument and NCE for PT-CID data. The best setting is Lumos@NCE=19. The grid search takes 114.2 seconds. (c) Fine-tuning for Sciex-6600 data using the best setting. (d) Fine-tuning for CID data using the best setting.

ProteomeTools also provides a lot of high-quality LIT-based CID data. Unlike beam-type CID including HCD, LIT-based CID cleaves peptides and detects fragment ions in the linear ion trap, it has some characteristics: (1) LIT-based fragmentation is a resonance-type fragmentation, under which most ions are fragmented only once (i.e. single cleavage); (2) LIT has a “1/3 rule”, meaning that low mass (m/z lower than 1/3 of the precursor m/z) can be hardly detected in the LIT; (3) LIT is a low-resolution mass analyzer. These three characteristics make CID spectra quite different from HCD spectra. Therefore, it is not surprising that the pre-trained model with RAW-file-recorded NCE=35 cannot give a good prediction for CID spectra (Figure 4a). With grid search, prediction by the pre-trained model with the best instrument and NCE values (Lumos@NCE=19) gets much better PCC_>0.90_ than that with Lumos@NCE=35. After fine-tuning by using 1000 CID spectra and 16 epochs with Lumos@NCE=19, the PCC_>0.90_ achieves 83.9% (Figure 4d).

Figure 5 clearly illustrates how much the prediction is improved for peptide “GFH-PDPEALK (3+)”. Predicted by the pre-trained model which is only trained by HCD data, the low-mass fragments get quite high intensities, but they could hardly get high intensities due to the “1/3 rule” in LIT-based CID (Figure 5a). After fine-tuning, the model learns the characteristics of LIT-based CID, and hence gives a better prediction (Figure 5c). Fine-tuning on LIT-based CID data with 1000 PSMs for 16 epochs costs less than 15 seconds.

**Figure 5:**
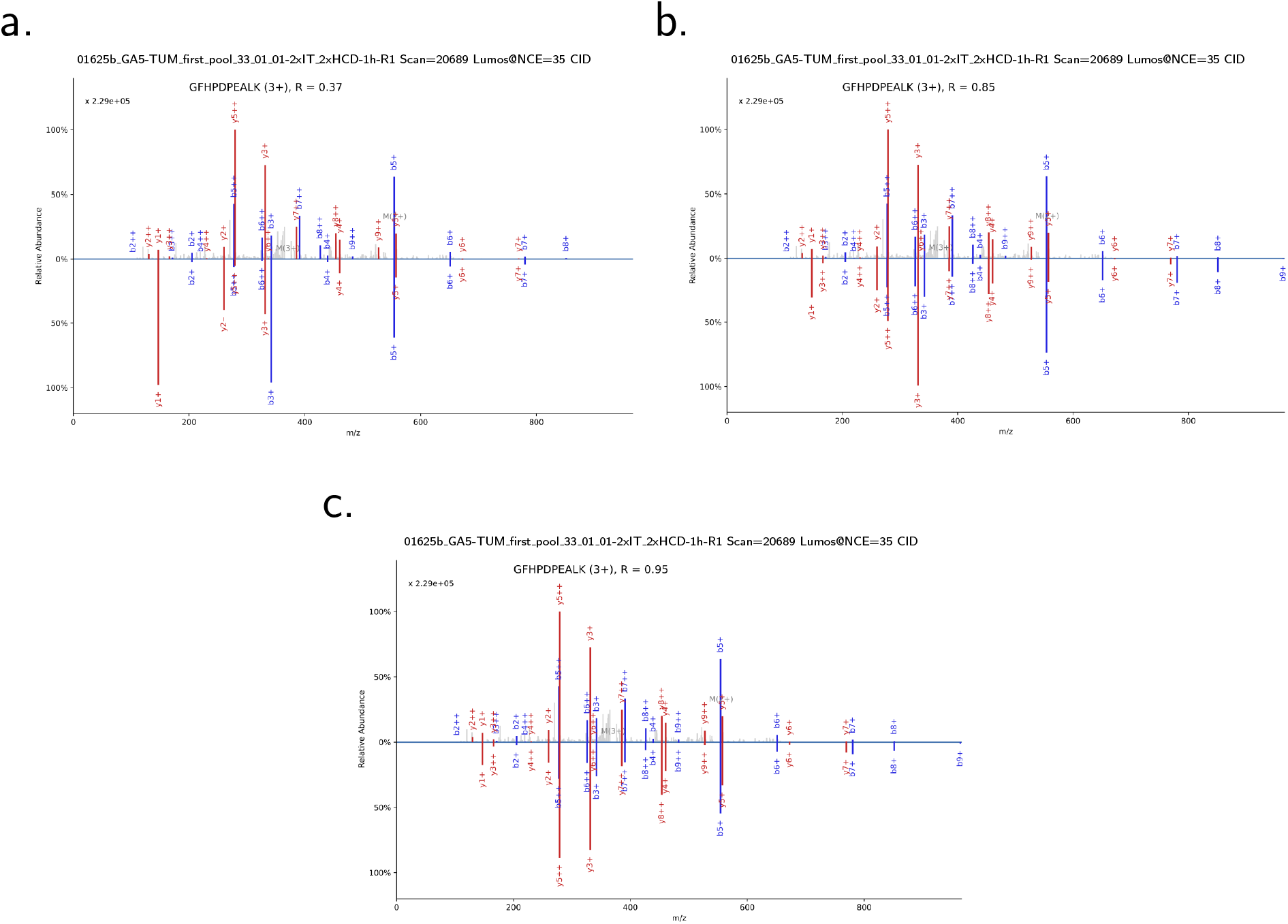
Examples of the prediction for the CID spectrum of peptide “GFHPDPEALK (3+)”. “R” refers the PCC. (a) Prediction with pre-trained model, Lumos@NCE=35. (b) Prediction with pre-trained model, Lumos@NCE=19. (c) Prediction with fine-tuned model, Lumos@NCE=19. The model is fine-tuned with 1000 PSMs and 16 epochs. The fine-tuning costs less than 15 seconds.

Therefore, few-shot learning coupled with grid search shows a great potential to train better models for unseen instrument types.

## Conclusions and Discussions

In this work, we show that a pre-trained pDeep model can be significantly improved using the few-shot learning technique. Few-shot learning aims to fine-tune the pDeep model with a small training set and a small number of training epochs and can increase the predicting accuracy within only seconds.

These years, Bruker’s timsTOF has attracted a lot of attention in proteomics.^28^ Although pDeep3 could not currently access the timsTOF data, we believe that fine-tuning could also be used for timsTOF data as it shows promising results for Sciex-6600 and LIT-based CID data.

Predicting the intensities of peptides with common or low-abundance PTMs has been discussed in pDeep2. It is also very promising that pDeep3 would be able to give a better prediction for peptides with PTMs. This would be very useful when users focus on PTM analysis.

In this work, all fine-tuning steps are done by CPU, hence we believe few-shot learning or transfer learning could be very useful to train a better model for users who do not have too many computational resources.

## Acknowledgement

The authors would like to thank Dr. Si-Min He for his invaluable supervision, our other col-leagues of pFind Team for their helpful discussions and suggestions, and Dr. Ruixiang Sun from National Institute of Biological Sciences for his advice about HCD and CID fragmentation techniques. This work was supported by the National Key Research and Development Program of China (No. 2016YFA0501301).

## Supporting Information Available

- **Note A:** Settings of the pre-trained model.
- **Note B:** Fine-tuning on HLA data.
- **Note C:** Building predicted spectral libraries for DIA analysis.
- **Data S1:** Test results of different combinations of *e* and *n* on all test datasets.
- **Data S2:** Intra-RAW and inter-RAW test results on all test datasets.

## Footnotes

* The pre-trained model includes improvments and bug fixes of the previous work ^9^ and is re-trained.

## Notes

http://pfind.ict.ac.cn/software/pdeep3

### Competing Interest Statement

The authors have declared no competing interest.

