## Supporting Information for "pDeep3: Towards More Accurate Spectrum Prediction with Fast Few-Shot Learning"

### Contents

**Note A: Settings of the pre-trained model** **S3**

**Note B: Fine-tuning on HLA data** **S4**

**Note C: Building predicted spectral libraries for DIA analysis** **S5**

**Data S1: Test results of different combinations of $e$ and $n$ on all test datasets** **S8**

**Data S2: Intra-RAW and inter-RAW test results on all test datasets** **S10**

**References** **S12**

### Note A: Settings of the pre-trained model

Table S1: Datasets used to train the pre-trained model

| Description | Lab | Instrument | NCE | #PSMs |
|---|---|---|---|---|
| mouse brain[S1] | Mann | QEHF | 27 | 221,106 |
| fission yeast[S2] | Mann | QE | 25 | 118,950 |
| HEK-293T[S3] | Gygi | QE | 25 | 244,177 |
| HPMa: adult CD4Tcells[S4] | Pandey | Elite | 32 | 77,901 |
| HPM: adult CD4Tcells | Pandey | Velos | 41 | 41,465 |
| HPM: adult CD4Tcells (gel) | Pandey | Velos | 41 | 56,172 |
| HPM: adult Lung | Pandey | Velos | 39 | 29,493 |
| ProteomeTools[S5] | Kuster | Lumos | 25 | 687,865 |
| ProteomeTools | Kuster | Lumos | 30 | 867,860 |
| ProteomeTools | Kuster | Lumos | 35 | 652,251 |

The data are searched using pFind3[S6] with the following settings.

- the corresponding SwissProt database of each species
- open-search mode
- precursor tolerance = $\pm$20 ppm
- fragment tolerance = $\pm$20 ppm
- number of missed cleavages $\leq 10$
- $6 \leq$ peptide lengths $\leq 100$

Only unmodified PSMs were kept at 0.01% FDR. For ProteomeTools, only peptides from the synthetic templates were kept.

#### Note B: Fine-tuning on HLA data

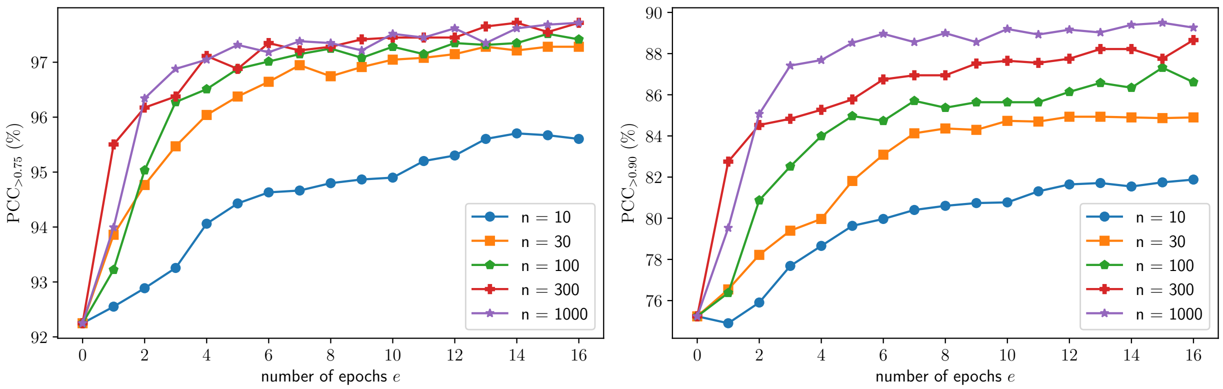

Figure S1: Performance on human leukocyte antigens (HLA) data, where $n$ is the number of spectra used for fine-tuning. The dataset, including search results, is available at ProteomeXchange repository (PXD004894),[S7] and two RAW files are used for fine-tuning and testing respectively. The pre-trained model is only trained with tryptic peptides, and, similarly, the figure shows that pDeep3 achieves significantly better performance on HLA data.

### Note C: Building predicted spectral libraries for DIA analysis

#### Spectral library generation

pDeep3 supports generating spectral libraries for EncyclopeDIA[S8] and OpenSWATH.[S9] For EncyclopeDIA analysis, pDeep3 generates SQLite-based .dlib library; for OpenSWATH analysis, pDeep3 generates SQLite-based .pqp library or text-based .tsv library. For the predicted libraries, pDeep3 predicts the retention time (RT) of the peptides using a pDeep3-build-in RT prediction model (pDeepRT, source codes can be found inside pDeep3 repository).

pDeep3 supports various formats of input peptide sequences, including the protein fasta files, spectral library files (.tsv, .dlib, etc.), pFind3 search results, MaxQuant search results, etc. When the input is a fasta file, pDeep3 automatically cleaves the protein sequences into peptide sequences and adds the user-specific fixed and variable modifications to the peptide sequences. The transfer learning technique used in pDeep2[S10] is also applicable in pDeep3 for modified peptides.

#### Test of OpenSWATH

DIA and DDA RAW data are downloaded from Chorus Project with ID 1105.[S11] The DDA RAW files (QEHF, NCE=27) are searched by pFind3 without the open search, and then the random selected 100 PSMs at 0.1% FDR are submitted to pDeep3 for fine-tuning with 2 epochs.

For DIA analysis, pan-human-library (PHL, in .tsv format) is downloaded from `https://db.systemsbiology.net/sbeams/cgi/downloadFile.cgi?name=phl004_canonical_s32_osw.csv;format=tsv;tmp_file=5dabd07a516696e7031d7e7cad45652a;raw_download=1`, and is converted to .pqp format by using "TargetedFileConverter" in OpenMS (version 2.4.0). We append decoy transitions by using "OpenSwathDecoyGenerator" with the "reverse" decoy method. Taking PHL as input, we generate the predicted .pqp PHL without fine-tuning ("pre-trained") and with fine-tuning ("tuned") based on the downloaded PHL, and the decoy method is set as "reverse" in pDeep3. OpenSWATH needs reference peptides (such as iRT peptides) for retention time (RT) alignment between RAW files and libraries, and hence we use peptides from endogenous proteins in human cells (actin and vinculin) in PHL for RT alignment. For the predicted libraries, we predict the RT of the library peptides as well as the reference peptides using pDeepRT. The DIA RAW files (QEHF, NCE=27) are converted to mzML by ProteoWizard[S12] with the default parameters. The searching parameters of OpenSWATH are: MS1 mass tolerance = ±10 ppm, MS2 mass tolerance = ±20 ppm, and "RT extraction window" = 600. The false discovery rate (FDR) is then estimated by pyprophet using the global-context estimation.[S13] The test results of PHL, "pre-trained" and "tuned" at 1% MS2 level FDR are shown in Figure S2a. The tuned PHL only identifies 98 more peptides than those from pre-trained PHL, because the pre-trained model already has very good prediction accuracy in the corresponding DDA RAW file (median PCCs are both 0.98 for pre-trained and tuned models).

#### Test of EncyclopeDIA

DIA and DDA RAW data are downloaded from PRIDE with ID PXD006722.[S14] The DDA RAW files (QEHF, NCE=27) are searched by pFind3 without open search, then the random selected 100 PSMs at 0.1% FDR are used for fine-tuning with 2 epochs.

For DIA search, PHL is downloaded from `https://bitbucket.org/searleb/encyclopedia/downloads/pan_human_library.dlib`, then the pre-trained and tuned PHL in .dlib format are generated for EncyclopeDIA. The DIA RAW files (QEHF, NCE=27) are converted to mzML, and then searched by EncyclopeDIA (version 0.9.0) using PHL, pre-trained PHL, and tuned PHL, respectively. The searching parameter "acquisition" is set as overlapping and other searching parameters are left as default. The searched results are demonstrated in Figure S2b. The median PCC is 0.95 for the pre-trained model, and increases to 0.96 after

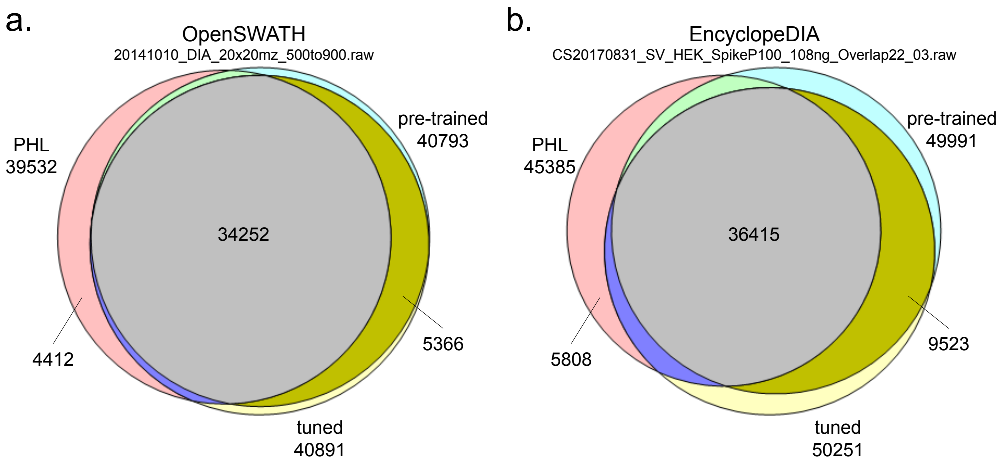

Figure S2: Identified peptides using OpenSWATH (a) and EncyclopeDIA (b) with PHL, predicted PHL without fine-tuning ("pre-trained"), and predicted PHL with fine-tuning ("tuned").

fine-tuning, resulting in 260 more identified peptides.

For both OpenSWATH and EncyclopeDIA, predicted (pre-trained and tuned) PHL can identify more peptides than the original library, demonstrating the potential advantages of spectrum prediction of pDeep3 for DIA analysis.

### Data S1: Test results of different combinations of $e$ and $n$ on all test datasets

PCC75

| dataset | size | #epoch | | | | | | | | |
|---|---|---|---|---|---|---|---|---|---|---|
| | | 0 | 1 | 2 | 3 | 4 | 5 | 6 | 7 | 8 |
| KV | 10 | 97.39% | 98.37% | 98.32% | 98.34% | 98.46% | 98.62% | 98.54% | 98.50% | 98.52% |
| | 30 | 97.39% | 98.37% | 98.64% | 98.54% | 98.57% | 98.45% | 98.54% | 98.68% | 98.62% |
| | 100 | 97.39% | 98.50% | 98.50% | 98.55% | 98.55% | 98.70% | 98.55% | 98.66% | 98.64% |
| | 300 | 97.39% | 98.50% | 98.46% | 98.39% | 98.46% | 98.41% | 98.54% | 98.46% | 98.46% |
| | 1000 | 97.39% | 98.55% | 98.48% | 98.68% | 98.55% | 98.68% | 98.61% | 98.59% | 98.66% |
| O2 | 10 | 96.23% | 96.46% | 96.61% | 96.81% | 96.85% | 96.85% | 96.88% | 96.92% | 96.97% |
| | 30 | 96.23% | 97.05% | 97.33% | 97.58% | 97.54% | 97.78% | 97.74% | 97.85% | 97.84% |
| | 100 | 96.23% | 97.52% | 97.93% | 97.86% | 97.86% | 97.96% | 97.88% | 97.92% | 98.02% |
| | 300 | 96.23% | 97.84% | 97.84% | 97.93% | 98.04% | 98.02% | 98.01% | 98.01% | 98.01% |
| | 1000 | 96.23% | 97.81% | 97.84% | 97.97% | 98.05% | 98.07% | 98.08% | 98.03% | 98.06% |
| OC | 10 | 92.96% | 93.44% | 94.16% | 94.09% | 94.16% | 94.22% | 94.06% | 94.20% | 94.48% |
| | 30 | 92.96% | 94.19% | 94.72% | 95.38% | 95.53% | 95.70% | 95.57% | 95.59% | 95.41% |
| | 100 | 92.96% | 95.26% | 95.96% | 96.68% | 96.80% | 96.80% | 96.88% | 96.85% | 96.83% |
| | 300 | 92.96% | 95.61% | 96.24% | 96.62% | 96.70% | 96.75% | 96.81% | 96.80% | 96.99% |
| | 1000 | 92.96% | 96.80% | 96.87% | 96.91% | 97.00% | 96.96% | 97.10% | 97.04% | 96.89% |
| OG | 10 | 91.61% | 92.33% | 93.40% | 93.82% | 93.85% | 93.55% | 93.55% | 93.62% | 93.85% |
| | 30 | 91.61% | 92.86% | 93.45% | 93.67% | 94.31% | 94.96% | 94.89% | 94.95% | 94.81% |
| | 100 | 91.61% | 94.88% | 95.54% | 95.65% | 95.51% | 95.22% | 95.40% | 95.55% | 95.62% |
| | 300 | 91.61% | 95.32% | 96.16% | 96.60% | 96.78% | 96.92% | 96.87% | 96.87% | 96.83% |
| | 1000 | 91.61% | 96.37% | 96.91% | 97.04% | 97.02% | 97.02% | 97.08% | 97.13% | 97.12% |
| OH | 10 | 94.45% | 94.86% | 95.21% | 95.34% | 95.35% | 95.48% | 95.43% | 95.21% | 95.24% |
| | 30 | 94.45% | 94.76% | 95.22% | 95.46% | 95.49% | 95.54% | 95.64% | 95.43% | 95.66% |
| | 100 | 94.45% | 95.59% | 96.01% | 95.90% | 95.99% | 95.95% | 95.99% | 95.87% | 95.94% |
| | 300 | 94.45% | 95.80% | 95.99% | 96.16% | 96.05% | 96.20% | 96.20% | 96.26% | 96.22% |
| | 1000 | 94.45% | 96.04% | 96.09% | 96.19% | 96.25% | 96.25% | 96.39% | 96.25% | 96.23% |
| OL | 10 | 94.23% | 94.35% | 94.42% | 94.67% | 94.92% | 95.04% | 95.15% | 95.01% | 95.09% |
| | 30 | 94.23% | 94.65% | 95.06% | 95.43% | 95.51% | 95.55% | 95.80% | 95.75% | 95.55% |
| | 100 | 94.23% | 95.25% | 96.07% | 96.29% | 96.53% | 96.36% | 96.63% | 96.53% | 96.63% |
| | 300 | 94.23% | 96.15% | 96.52% | 96.57% | 96.70% | 96.59% | 96.63% | 96.75% | 96.49% |
| | 1000 | 94.23% | 95.77% | 96.45% | 96.76% | 96.71% | 96.78% | 96.74% | 96.82% | 96.79% |
| PV | 10 | 83.72% | 92.45% | 94.75% | 95.22% | 95.58% | 95.71% | 95.51% | 95.47% | 95.48% |
| | 30 | 83.72% | 95.01% | 95.81% | 95.69% | 96.00% | 95.72% | 95.96% | 95.69% | 96.06% |
| | 100 | 83.72% | 95.99% | 96.25% | 96.21% | 96.32% | 96.42% | 96.54% | 96.40% | 96.52% |
| | 300 | 83.72% | 96.17% | 96.38% | 96.53% | 96.40% | 96.60% | 96.63% | 96.68% | 96.56% |
| | 1000 | 83.72% | 95.90% | 96.31% | 96.25% | 96.43% | 96.52% | 96.47% | 96.68% | 96.64% |
| PL | 10 | 99.51% | 99.51% | 99.51% | 99.49% | 99.49% | 99.49% | 99.49% | 99.49% | 99.49% |
| | 30 | 99.51% | 99.48% | 99.48% | 99.48% | 99.48% | 99.48% | 99.49% | 99.49% | 99.48% |
| | 100 | 99.51% | 99.51% | 99.48% | 99.48% | 99.48% | 99.48% | 99.48% | 99.48% | 99.48% |
| | 300 | 99.51% | 99.49% | 99.49% | 99.49% | 99.49% | 99.48% | 99.48% | 99.48% | 99.49% |
| | 1000 | 99.51% | 99.49% | 99.51% | 99.51% | 99.51% | 99.51% | 99.51% | 99.51% | 99.51% |

PCC90

| dataset | size | #epoch | | | | | | | | |
|---|---|---|---|---|---|---|---|---|---|---|
| | | 0 | 1 | 2 | 3 | 4 | 5 | 6 | 7 | 8 |
| KV | 10 | 80.64% | 89.23% | 90.37% | 90.53% | 90.78% | 90.96% | 91.12% | 91.05% | 91.01% |
| | 30 | 80.64% | 90.34% | 91.28% | 91.00% | 91.55% | 91.14% | 91.32% | 91.68% | 91.80% |
| | 100 | 80.64% | 91.48% | 91.84% | 92.16% | 91.98% | 92.34% | 92.35% | 92.27% | 92.25% |
| | 300 | 80.64% | 92.02% | 91.89% | 92.10% | 92.19% | 92.52% | 92.64% | 92.41% | 92.52% |
| | 1000 | 80.64% | 92.30% | 92.23% | 92.62% | 92.77% | 93.02% | 93.16% | 92.98% | 93.23% |
| O2 | 10 | 81.29% | 83.01% | 83.98% | 84.43% | 84.42% | 84.59% | 84.77% | 84.86% | 84.79% |
| | 30 | 81.29% | 85.86% | 87.34% | 87.58% | 88.27% | 89.29% | 89.19% | 89.45% | 89.66% |
| | 100 | 81.29% | 87.95% | 89.78% | 89.76% | 90.22% | 90.31% | 90.46% | 90.72% | 91.05% |
| | 300 | 81.29% | 89.82% | 90.60% | 90.92% | 91.21% | 91.18% | 91.20% | 91.38% | 91.15% |
| | 1000 | 81.29% | 89.93% | 90.73% | 91.35% | 91.45% | 91.57% | 91.60% | 91.48% | 91.46% |
| OC | 10 | 73.41% | 72.82% | 74.27% | 74.85% | 76.21% | 76.68% | 76.31% | 76.70% | 76.94% |
| | 30 | 73.41% | 75.36% | 76.32% | 78.85% | 78.91% | 80.13% | 79.79% | 79.19% | 79.02% |
| | 100 | 73.41% | 79.90% | 81.58% | 82.87% | 83.61% | 83.57% | 83.63% | 83.82% | 83.48% |
| | 300 | 73.41% | 80.77% | 82.40% | 83.39% | 83.97% | 83.48% | 84.45% | 83.98% | 84.59% |
| | 1000 | 73.41% | 84.66% | 84.99% | 85.56% | 85.62% | 85.66% | 85.90% | 85.47% | 85.80% |
| OG | 10 | 66.87% | 69.61% | 72.51% | 72.63% | 72.87% | 72.47% | 72.55% | 71.71% | 72.69% |
| | 30 | 66.87% | 72.17% | 73.93% | 74.76% | 76.94% | 77.56% | 76.97% | 77.79% | 77.58% |
| | 100 | 66.87% | 77.05% | 79.28% | 79.64% | 80.30% | 78.97% | 79.86% | 79.59% | 79.12% |
| | 300 | 66.87% | 77.42% | 80.53% | 82.27% | 82.20% | 82.51% | 82.40% | 82.50% | 82.54% |
| | 1000 | 66.87% | 81.11% | 83.25% | 83.63% | 83.74% | 83.16% | 83.78% | 84.02% | 84.06% |
| OH | 10 | 80.47% | 82.27% | 83.89% | 84.37% | 84.44% | 83.62% | 83.44% | 84.09% | 84.24% |
| | 30 | 80.47% | 82.80% | 84.95% | 85.74% | 85.62% | 85.77% | 86.23% | 85.47% | 86.41% |
| | 100 | 80.47% | 86.02% | 88.03% | 87.78% | 88.08% | 88.32% | 88.03% | 88.09% | 88.26% |
| | 300 | 80.47% | 87.65% | 88.16% | 88.72% | 88.43% | 88.86% | 88.59% | 88.86% | 88.69% |
| | 1000 | 80.47% | 88.44% | 88.85% | 88.94% | 89.18% | 89.19% | 89.22% | 89.04% | 89.07% |
| OL | 10 | 75.28% | 74.70% | 76.44% | 77.07% | 77.71% | 78.03% | 78.80% | 78.58% | 78.96% |
| | 30 | 75.28% | 77.66% | 80.00% | 81.72% | 82.13% | 82.59% | 82.67% | 82.68% | 82.41% |
| | 100 | 75.28% | 80.85% | 83.62% | 84.67% | 85.37% | 84.91% | 85.88% | 85.56% | 85.87% |
| | 300 | 75.28% | 83.76% | 85.70% | 85.85% | 86.07% | 85.93% | 86.09% | 86.35% | 85.47% |
| | 1000 | 75.28% | 83.27% | 85.87% | 86.44% | 86.31% | 86.51% | 86.57% | 86.45% | 86.70% |
| PV | 10 | 48.66% | 66.43% | 77.83% | 82.20% | 84.28% | 84.49% | 84.21% | 84.52% | 84.98% |
| | 30 | 48.66% | 82.61% | 86.39% | 86.47% | 87.43% | 87.42% | 87.79% | 87.04% | 87.79% |
| | 100 | 48.66% | 87.18% | 88.10% | 88.06% | 88.64% | 88.82% | 88.99% | 88.64% | 88.99% |
| | 300 | 48.66% | 88.02% | 88.49% | 88.91% | 88.50% | 89.13% | 89.13% | 89.31% | 88.96% |
| | 1000 | 48.66% | 87.86% | 88.78% | 88.48% | 88.98% | 89.27% | 89.16% | 89.48% | 89.41% |
| PL | 10 | 97.05% | 97.08% | 97.14% | 97.12% | 97.11% | 97.12% | 97.11% | 97.09% | 97.09% |
| | 30 | 97.05% | 97.03% | 96.91% | 96.99% | 97.09% | 97.08% | 97.06% | 97.06% | 97.00% |
| | 100 | 97.05% | 97.14% | 97.12% | 97.12% | 97.15% | 97.18% | 97.08% | 97.08% | 97.05% |
| | 300 | 97.05% | 97.08% | 97.06% | 97.11% | 97.12% | 97.09% | 97.08% | 97.14% | 97.15% |
| | 1000 | 97.05% | 97.03% | 97.08% | 97.03% | 97.09% | 97.11% | 97.12% | 97.14% | 97.02% |

### Data S2: Intra-RAW and inter-RAW test results on all test datasets

PCC75

| KV | ts-1 | ts-2 | ts-3 | ts-4 | ts-5 | | σ= |
|---|---|---|---|---|---|---|---|
| tr-1 | 98.65% | 98.34% | 98.66% | 97.40% | 99.21% | | 0.56% |
| tr-2 | 98.76% | 98.69% | 98.75% | 97.74% | 99.27% | | |
| tr-3 | 98.65% | 98.42% | 98.66% | 97.57% | 99.23% | | |
| tr-4 | 98.78% | 98.53% | 98.73% | 97.62% | 99.21% | | |
| tr-5 | 98.57% | 98.48% | 98.66% | 97.50% | 99.21% | | |
| O2 | ts-1 | ts-2 | ts-3 | ts-4 | ts-5 | | σ= |
| tr-1 | 97.64% | 97.89% | 97.54% | 97.92% | 97.89% | | 0.17% |
| tr-2 | 97.74% | 98.00% | 97.61% | 97.96% | 98.01% | | |
| tr-3 | 97.74% | 97.98% | 97.70% | 98.01% | 97.99% | | |
| tr-4 | 97.50% | 97.85% | 97.55% | 97.90% | 97.89% | | |
| tr-5 | 97.62% | 97.88% | 97.56% | 97.95% | 97.91% | | |
| OC | ts-1 | ts-2 | ts-3 | ts-4 | ts-5 | | σ= |
| tr-1 | 96.90% | 96.46% | 96.40% | 96.55% | 97.20% | | 0.39% |
| tr-2 | 96.67% | 95.88% | 95.83% | 96.21% | 96.54% | | |
| tr-3 | 96.91% | 96.42% | 96.34% | 96.66% | 97.24% | | |
| tr-4 | 96.64% | 96.31% | 96.01% | 96.34% | 96.87% | | |
| tr-5 | 96.41% | 95.87% | 95.88% | 96.16% | 96.63% | | |
| OG | ts-1 | ts-2 | ts-3 | ts-4 | ts-5 | | σ= |
| tr-1 | 96.09% | 95.98% | 96.29% | 97.00% | 96.18% | | 0.95% |
| tr-2 | 95.54% | 95.81% | 95.94% | 96.67% | 95.63% | | |
| tr-3 | 94.79% | 94.76% | 95.19% | 96.02% | 94.97% | | |
| tr-4 | 93.78% | 93.77% | 93.56% | 94.75% | 93.81% | | |
| tr-5 | 94.83% | 94.47% | 94.72% | 95.89% | 94.62% | | |
| OH | ts-1 | ts-2 | ts-3 | ts-4 | ts-5 | | σ= |
| tr-1 | 95.50% | 96.02% | 98.09% | 98.02% | 97.04% | | 1.00% |
| tr-2 | 95.48% | 95.69% | 97.58% | 97.65% | 96.59% | | |
| tr-3 | 95.51% | 95.82% | 97.78% | 97.77% | 96.85% | | |
| tr-4 | 95.59% | 95.90% | 97.87% | 97.93% | 96.90% | | |
| tr-5 | 95.35% | 95.63% | 97.68% | 97.70% | 96.70% | | |
| OL | ts-1 | ts-2 | ts-3 | ts-4 | ts-5 | | σ= |
| tr-1 | 96.57% | 96.23% | 95.50% | 96.25% | 98.61% | | 1.18% |
| tr-2 | 96.15% | 95.67% | 94.76% | 95.23% | 98.31% | | |
| tr-3 | 96.32% | 95.51% | 95.10% | 95.83% | 98.39% | | |
| tr-4 | 96.40% | 95.96% | 95.45% | 96.16% | 98.52% | | |
| tr-5 | 96.08% | 95.59% | 95.00% | 95.64% | 98.41% | | |
| PV | ts-1 | ts-2 | ts-3 | ts-4 | ts-5 | | σ= |
| tr-1 | 96.84% | 96.45% | 96.42% | 96.45% | 97.36% | | 0.35% |
| tr-2 | 96.51% | 96.14% | 96.24% | 96.16% | 96.88% | | |
| tr-3 | 97.03% | 96.65% | 96.63% | 96.69% | 97.25% | | |
| tr-4 | 96.67% | 96.23% | 96.31% | 96.27% | 96.93% | | |
| tr-5 | 96.75% | 96.25% | 96.32% | 96.36% | 97.12% | | |
| PL | ts-1 | ts-2 | ts-3 | ts-4 | ts-5 | | σ= |
| tr-1 | 99.40% | 99.46% | 99.09% | 98.48% | 98.52% | | 0.47% |
| tr-2 | 99.24% | 99.48% | 98.99% | 98.35% | 98.26% | | |
| tr-3 | 99.26% | 99.49% | 98.97% | 98.37% | 98.29% | | |
| tr-4 | 99.33% | 99.49% | 99.13% | 98.41% | 98.44% | | |
| tr-5 | 99.34% | 99.26% | 99.15% | 98.13% | 98.97% | | |

PCC90

| KV | ts-1 | ts-2 | ts-3 | ts-4 | ts-5 | | σ= |
|---|---|---|---|---|---|---|---|
| tr-1 | 93.21% | 91.84% | 94.26% | 92.69% | 95.84% | | 1.51% |
| tr-2 | 93.10% | 91.65% | 93.32% | 92.46% | 96.10% | | |
| tr-3 | 93.25% | 91.74% | 93.75% | 92.57% | 95.98% | | |
| tr-4 | 94.25% | 92.17% | 93.89% | 93.06% | 96.22% | | |
| tr-5 | 92.88% | 91.11% | 93.05% | 92.19% | 95.86% | | |
| O2 | ts-1 | ts-2 | ts-3 | ts-4 | ts-5 | | σ= |
| tr-1 | 89.70% | 90.12% | 89.40% | 89.34% | 89.24% | | 0.38% |
| tr-2 | 90.04% | 90.44% | 89.52% | 89.73% | 89.91% | | |
| tr-3 | 89.92% | 90.14% | 89.97% | 89.57% | 89.78% | | |
| tr-4 | 90.00% | 90.43% | 89.69% | 90.27% | 90.13% | | |
| tr-5 | 89.27% | 89.50% | 89.07% | 89.40% | 89.66% | | |
| OC | ts-1 | ts-2 | ts-3 | ts-4 | ts-5 | | σ= |
| tr-1 | 82.59% | 82.43% | 81.83% | 80.58% | 82.98% | | 1.14% |
| tr-2 | 82.02% | 81.50% | 80.87% | 80.15% | 83.01% | | |
| tr-3 | 82.71% | 82.98% | 82.21% | 81.61% | 84.34% | | |
| tr-4 | 82.71% | 82.65% | 81.45% | 81.13% | 83.26% | | |
| tr-5 | 81.03% | 80.60% | 80.41% | 79.67% | 82.51% | | |
| OG | ts-1 | ts-2 | ts-3 | ts-4 | ts-5 | | σ= |
| tr-1 | 79.91% | 79.75% | 80.28% | 81.47% | 81.13% | | 1.90% |
| tr-2 | 77.86% | 78.05% | 78.42% | 79.97% | 79.89% | | |
| tr-3 | 76.78% | 77.34% | 78.62% | 79.74% | 78.86% | | |
| tr-4 | 73.70% | 75.26% | 75.72% | 76.96% | 76.40% | | |
| tr-5 | 78.05% | 78.44% | 78.91% | 80.36% | 79.79% | | |
| OH | ts-1 | ts-2 | ts-3 | ts-4 | ts-5 | | σ= |
| tr-1 | 87.17% | 87.92% | 90.15% | 89.47% | 88.58% | | 1.19% |
| tr-2 | 85.72% | 86.31% | 87.71% | 87.79% | 86.83% | | |
| tr-3 | 86.70% | 87.35% | 89.23% | 89.05% | 87.91% | | |
| tr-4 | 87.13% | 87.78% | 89.84% | 89.57% | 88.36% | | |
| tr-5 | 86.26% | 86.85% | 88.24% | 88.64% | 87.47% | | |
| OL | ts-1 | ts-2 | ts-3 | ts-4 | ts-5 | | σ= |
| tr-1 | 85.26% | 84.24% | 83.46% | 84.59% | 92.38% | | 3.73% |
| tr-2 | 82.13% | 81.70% | 81.04% | 80.29% | 90.53% | | |
| tr-3 | 83.33% | 81.99% | 81.15% | 82.29% | 91.08% | | |
| tr-4 | 83.48% | 83.14% | 82.46% | 83.30% | 91.69% | | |
| tr-5 | 83.90% | 82.77% | 82.13% | 83.24% | 91.77% | | |
| PV | ts-1 | ts-2 | ts-3 | ts-4 | ts-5 | | σ= |
| tr-1 | 89.99% | 88.39% | 89.14% | 89.48% | 90.56% | | 0.76% |
| tr-2 | 89.78% | 88.30% | 89.12% | 89.37% | 90.80% | | |
| tr-3 | 90.23% | 88.62% | 89.09% | 89.45% | 90.39% | | |
| tr-4 | 89.85% | 88.31% | 88.75% | 89.25% | 90.49% | | |
| tr-5 | 90.02% | 88.45% | 89.10% | 89.32% | 90.36% | | |
| PL | ts-1 | ts-2 | ts-3 | ts-4 | ts-5 | | σ= |
| tr-1 | 96.26% | 97.13% | 95.14% | 93.85% | 87.10% | | 3.77% |
| tr-2 | 96.13% | 97.25% | 94.95% | 93.69% | 85.78% | | |
| tr-3 | 96.13% | 97.03% | 95.06% | 93.75% | 86.22% | | |
| tr-4 | 96.24% | 97.20% | 95.06% | 93.83% | 87.22% | | |
| tr-5 | 96.09% | 86.81% | 94.55% | 91.72% | 93.69% | | |
